# Single cell profiling of γδ hepatosplenic T-cell lymphoma unravels tumor cell heterogeneity associated with disease progression

**DOI:** 10.1101/2022.01.17.476575

**Authors:** Wei Song, Haixi Zhang, Fan Yang, Kiichi Nakahira, Cheng Wang, Keqian Shi, Ruoyu Zhang

## Abstract

Hepatosplenic T-cell lymphoma (HSTCL), mostly derived from γδ T cells, is a rare but very aggressive lymphoma with poor outcomes. The molecular pathogenesis driving HSTCL is largely unknown while only limited treatment options are available. In this study, by performing paired single cell RNA-seq and T cell receptor (TCR) sequencing on biopsies from a HSTCL patient pre- and post-chemotherapy treatments, we characterized unique gene expressing signatures of malignant γδ T cells, with a set of marker genes were newly identified in HSTCL (AREG, PLEKHA5, VCAM1 etc.). Although the malignant γδ T cells were expanded from a single TCR clonotype according to their TCR identities, they evolved into two transcriptional distinct tumor subtypes during the disease progression. The Tumor_1 subtype was dominant in pre-treatment samples with highly aggressive phenotypes. While the Tumor_2 had relative mild cancer hallmark signatures but expressed genes associated with tumor survival signal and drug resistance (IL32, TOX2, AIF1, AKAP12 etc.), and finally became the main tumor subtype post-treatment. We further dissected the tumor microenvironment of the HSTCL and noticed that CD8 memory T cells were clonal expanded post-treatment. In addition, we discovered dynamically rewiring cell-cell interaction networks during the treatment. The tumor cells had reduced communications with the microenvironment post-treatment. Our study reveals heterogenous and dynamic tumor and microenvironment underlying pathogenesis of HSTCL and may contribute to identify novel targets for diagnosis and cure of HSTCL in the future.

## Introduction

Hepatosplenic T-cell lymphoma (HSTCL) is a rare subtype of peripheral T-cell lymphoma (PTCL), mostly derived from γδ T cells. In a recent report, HSTCL accounts for 2% of all T-cell lymphoma subtypes worldwide^1^. In earlier surveys, it accounts 3% in the United States, 2.3% in Europe, and 0.2% in Asia^2^. The clinical presentations of HSTCL is highly aggressive and usually with a fatal outcome. The median overall survival time is only 1-2 years^3–6^ after diagnosis. Because it is rare, no treatments have become established as internationally recognized standards of care. Clinical trials are also unusual. Different combinations of chemotherapies have been used to treat HSTCL. In the earliest patient series, CHOP or CHOP-like regimens were reported as the most frequently used for induction treatment. More intensive chemotherapy regimens have been used as well, such as Hyper-CVAD or Hyper-CVAD-like regimens, and ICE or IVAC regiments^3^. However, no single regimen is clearly superior^3–6^. In addition, relapses occur often and early. In one series, the median time to relapse among patients achieving a complete response to induction therapy with CHOP or CHOP-like regimens was 3–16 months^3^. The efficient treatment modalities for HSTCL remain to be defined.

Given the aggressive, fatal nature of HSTCL, there is a critical need to identify putative molecular targets and develop novel therapeutic approaches. However, the pathogenesis of HSTCL remains largely unknown. Only a few molecular studies were conducted for this disease. Travert et al reported a distinct gene-expression profile of HSTCL, differentiating it from other T-cell lymphomas, and provided rationale for exploring new therapeutic options such as Syk inhibitors and demethylating agents^7^. Ferreiro et al integrated transcriptomic and genomic analysis to demonstrate that chromosome 7 imbalances were the driver events in HSTCL and identified a set of genes, marking HSTCL from other malignancies^8^.

To overcome the knowledge gap, in this study, we presented the first single cell analysis for HSTCL. We collected bone marrow and PBMC biopsies pre- and post-treatment from a 17-year-old female patient, and performed paired single cell RNA-seq (scRNA-seq) and T cell receptor sequencing (scTCR-seq). We established a cell types landscape for HSTCL and characterized the molecular signature of malignant γδ T cells from this rare lymphoma. We further revealed heterogeneities among the tumor cells and identified a tumor subset might be associated with disease progression and drug resistance. We also reconstructed the tumor microenvironment (TME) of HSTCL, and investigated how the TME were interacting with tumor cells and reforming under therapeutic conditions.

## Methods

### Patient recruitment and ethics statement

This study was approved by the Ethics Committee of The First People’s Hospital of Yunnan Province, China. The recruited patient gave informed consent at hospitalization.

### Clinical examinations

Computed tomography (CT) scanning was performed to assess the potential pulmonary infection and status of lymph nodes, liver and spleen. The scanning was ranged from the foramen magnum to the inferior margin of the symphysis pubis. Raw CT data was reconstructed with 1mm thickness to estimate liver and spleen volumes using the Post-processing workstation (Syngo via, Siemens, Germany).

In each hospitalization, peripheral blood and bone marrow samples were collected for smear cytology using Wright staining. The staining was performed with following steps: 1) Add Wright-Giemsa Solution A (~ 0.5~0.8ml) to the smear and stain for 1 min. 2) Add Wright-Giemsa Solution B (2~3 volumes of Solution A) onto Solution A and mix thoroughly then stain for 5~10mins. 3) Rinse with water gently, dry and examine the slide using a microscope.

The antibodies used in the flow cytometry analysis were CD10, CD16, CD138, CD45, CD56, CD2, CD3, HLA-DR, CD34, CD4, CD8, CD19, CD5, CD117, CD7, TCRαβ, TCRγδ, CD38, CD57, CD64 (BD Bioscience, US). The cells were washed in phosphate-buffered saline (PBS) and stained with a cocktail of cell surface antibodies for 20 min. Lysing solution (BD Bioscience, US) was then added to remove red blood cells. The cells were then washed and resuspended in PBS and analyzed by flow cytometry (FACS Canto, Bioscience, US).

TCR rearrangements were detected by IdentiClone® TCRB + TCRG Gene Clonality Assay following the manufacturer’s instructions (Invivoscribe, San Diego, US)

### Single cells RNA and TCR sequencing

Mononuclear cells were isolated from whole blood and bone marrow by centrifugation and resuspended with freezing medium. The cells were then frozen in a freezing container in a −80°C freezer. On the date of experiment, the cells were thawed using a water bath at 37°C and loaded into Chromium microfluidic chips and barcoded within a 10X Chromium Controller (10X Genomics, US). For transcriptome, procedures were performed with reagents: Chromium Next GEM Single Cell 5’ Library & Gel Bead Kit v1.1 (10X Genomics, Cat. No. 1000165). TCR enrichment was carried out using the Chromium Single Cell V(D)J Enrichment Kit, Human T Cell (10X Genomics, Cat. No. 1000005) for αβ transcripts, or customer primers for γδ TCR transcripts9. All the libraries were sequenced in a PE150 mode (Pair-End for 150bp read) on the NovaSeq 6000 platform (Illumina, US).

### Single cell data analysis

Raw sequencing data were processed by Cell Ranger version 6.0.2 (10X Genomics) to generate gene expression matrix and assemble TCR sequences with human GRCh38 reference genome. scRNA-seq data analysis were performed using Scanpy^10^. In quality control step, cells were filtered by the following criteria: 1) mitochondrial abundance < 10%, 2) minimum gene detected > 400, 3) Maximum UMI < 35000. Potential doublets were detected and removed by Scrublet^11^ and manual inspections. Principal component analysis (PCA) was applied on the highly variable genes and cells were projected and visualized in 2D dimensions using uniform manifold approximation and projection (UMAP) on the first 30 PCs. Cells were clustered by Leiden algorithm implemented in Scanpy. Differentially expressed genes were identified using a cut-off of |Fold Change| > 2, adjusted p value < 0.05 and gene expressed > 10% in the higher expression group. Gene enrichment analysis was performed using R package clusterProfiler^12^. TCR data was analyzed with Scirpy^13^. R package InferCNV was used to infer copy number variation (https://github.com/broadinstitute/infercnv) with following parameters (cut-off=0.1, cluster_by_groups=TRUE, denoise=TRUE, HMM=TRUE, analysis_mode=’subclusters’). CellChat R package^14^ was applied to identify cell-cell interactions.

### Data sharing statement

The sequencing data was deposited at Gene Expression Omnibus (GEO) under accession no. GSE193220.

## Results

### Clinical characteristics of the studied patient with HSTCL

The detailed clinical characteristics of the recruited patient with HSTCL were summarized in **Table. 1**. This 17-years old female patient was diagnosed γδ HSTCL on May 18 2021 (**Fig. 1A**), initially presented with fever and fatigue, and subsequently developed severe hemorrhage due to thrombocytopenia (**Table. 1**). CT scanning revealed splenomegaly and hepatomegaly (estimated spleen volume: 910.56 cm^3^ and liver volume: 1617.21 cm^3^ **Fig. 1B** left), while lymphadenopathy was absent (**Fig. S1**). Neoplastic lymphocytes were identified from bone marrow aspirate (**Fig. 1C**), which were large, round or irregular cells, with abundant cytoplasm but no or few particles in cytoplasm. The nuclei were round or irregular, depressed and distorted, with dense chromatin and 1-3 inconspicuous nucleoli, PAS glycogen staining was positive. The abnormal cells accounted for ~50% of lymphocytes. By flow cytometry, the neoplastic lymphocytes were found to be CD2+, CD3+, CD16+, CD45+, TCRγ/δ+, CD7P+, CD4-, CD8- and TCR α/β-, which were the typical immunophenotypes of γδ HSTCL^4,6^ **Fig. 1D**). TCR rearrangement assay further confirmed clonal rearrangement of TCR β (TCRB) and γ (TCRG) genes.

**Table 1.**
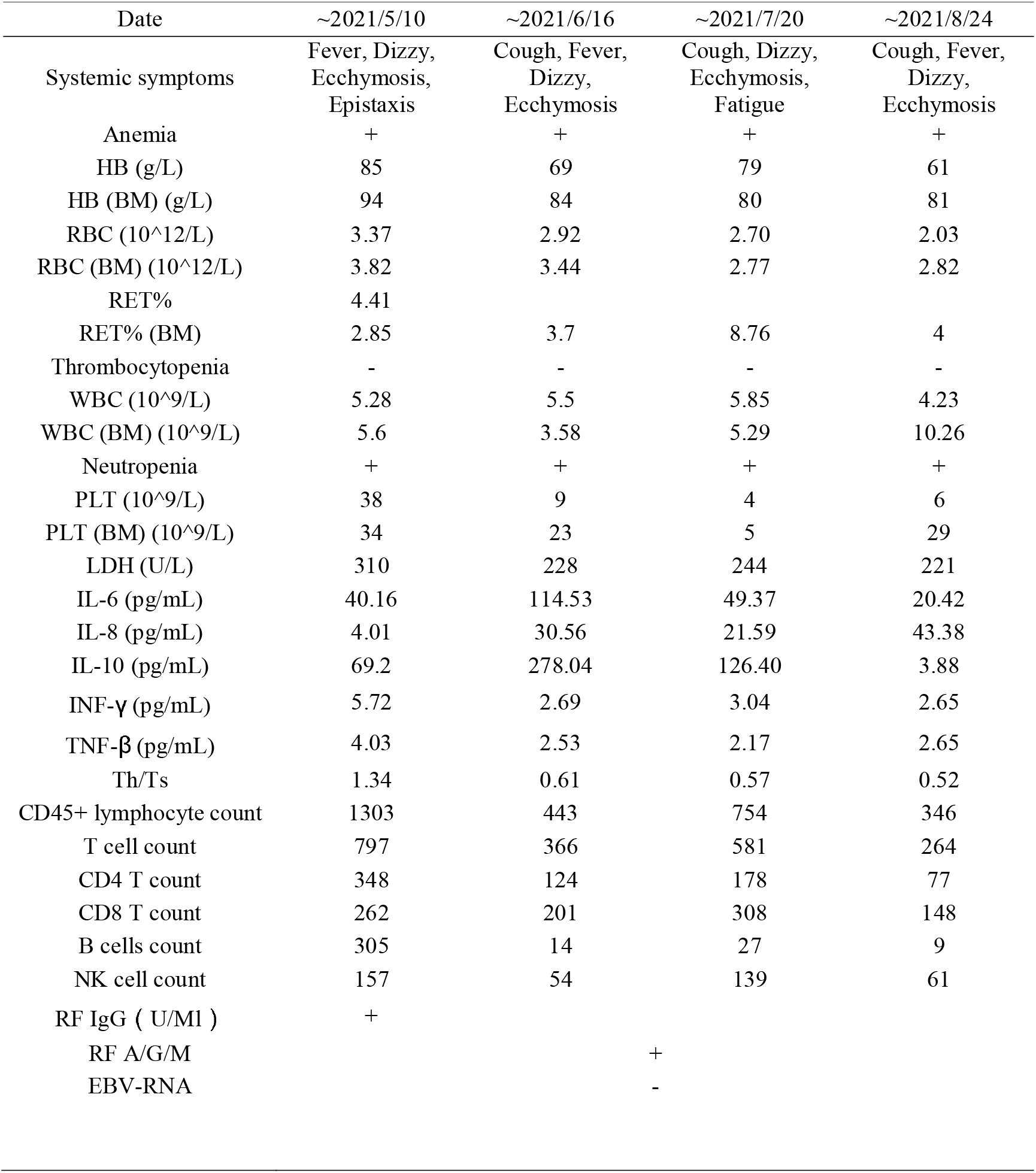
The clinical characteristics of the studied patient

**Figure 1.**
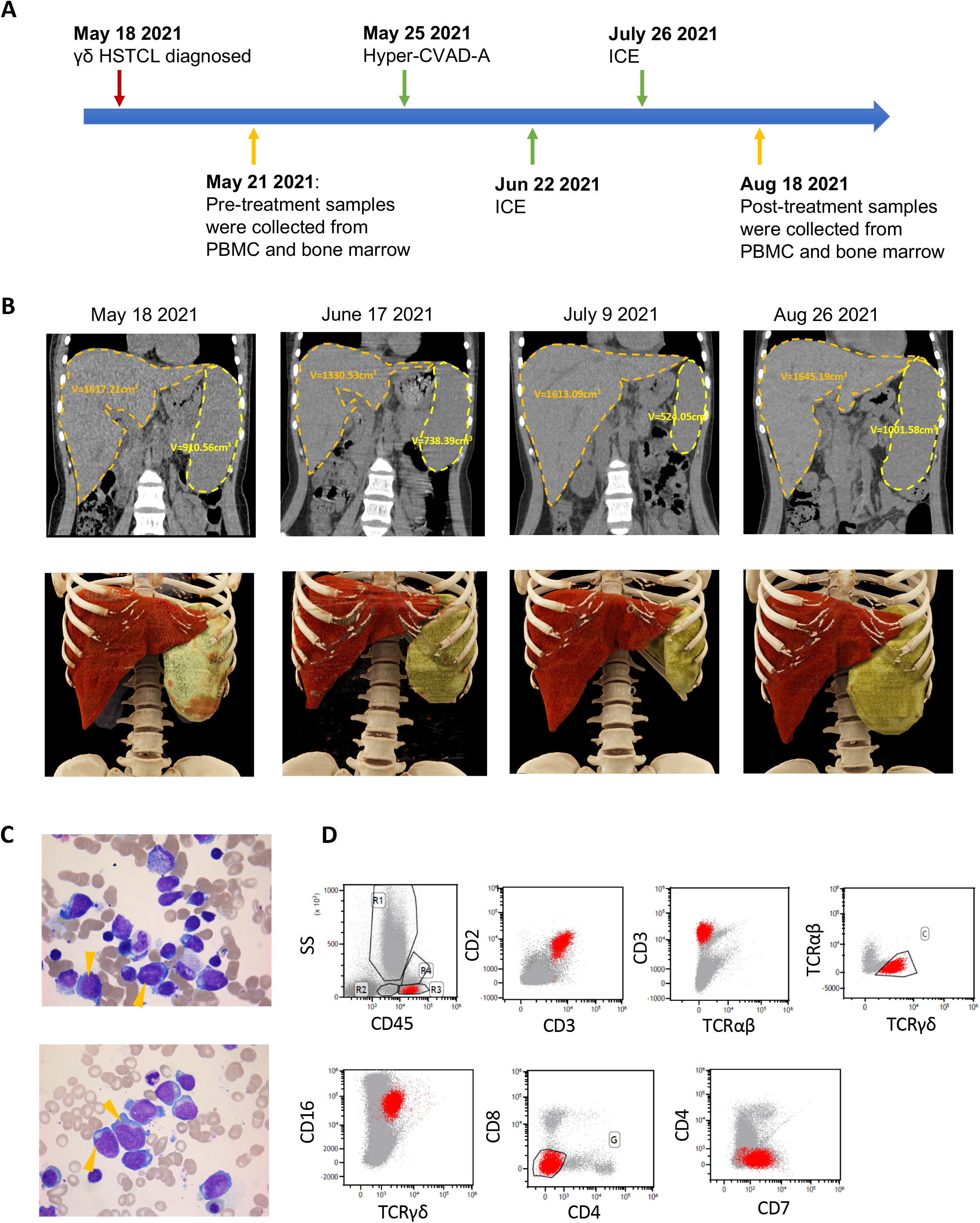
Clinical presentations of the studied HSTCL patient. (A) Timeline of the patient diagnosis, chemotherapy administration and biopsies collected for single cell sequencing. (B) CT scan and 3D image reconstruction during the disease progression, the volume of the spleen and liver was measured and labeled. (C) Wright-staining of bone marrow aspirate, arrows indicate selected typical neoplastic T cells. (D) Immunophenotypes of the neoplastic γδ T cells identified by flow cytometry.

After diagnosis, the patient began to receive chemotherapy hyper-CVAD-A regimen as induction treatment (**Fig. 1A, Table. 2**). During the first course of chemotherapy, the patient’s body temperature returned to the normal and the overall conditions was relatively stable. Just 17 days after the first chemotherapy completed, the patient was hospitalized again due to fever and thrombocytopenia. The patient was then received ICE regimen (**Fig. 1A Table. 2**), which achieved similar therapeutic outcomes as hyper-CVAD-A. Separated by 32 days, another course of ICE regimen was administrated to the patient (**Fig. 1A Table. 2)**. During each chemotherapy period, the indexes of routine blood examinations such as red blood cell count, white blood cell count, platelet count could gradually return to the normal range. However, the symptoms relapsed soon after the treatments were completed. During the second course of chemotherapy, CT examination revealed significantly reduced spleen volume (524.05 cm^3^) while liver volume stayed similar size (1613.09 cm^3^, **Fig. 1B** third column), but after the third course of treatment, the spleen and liver enlarged again to 1001.58 cm^3^ and 1645.19 cm^3^ (**Fig. 1B**. right). These results suggested HSTCL was very aggressive and refractory.

**Table 2.**
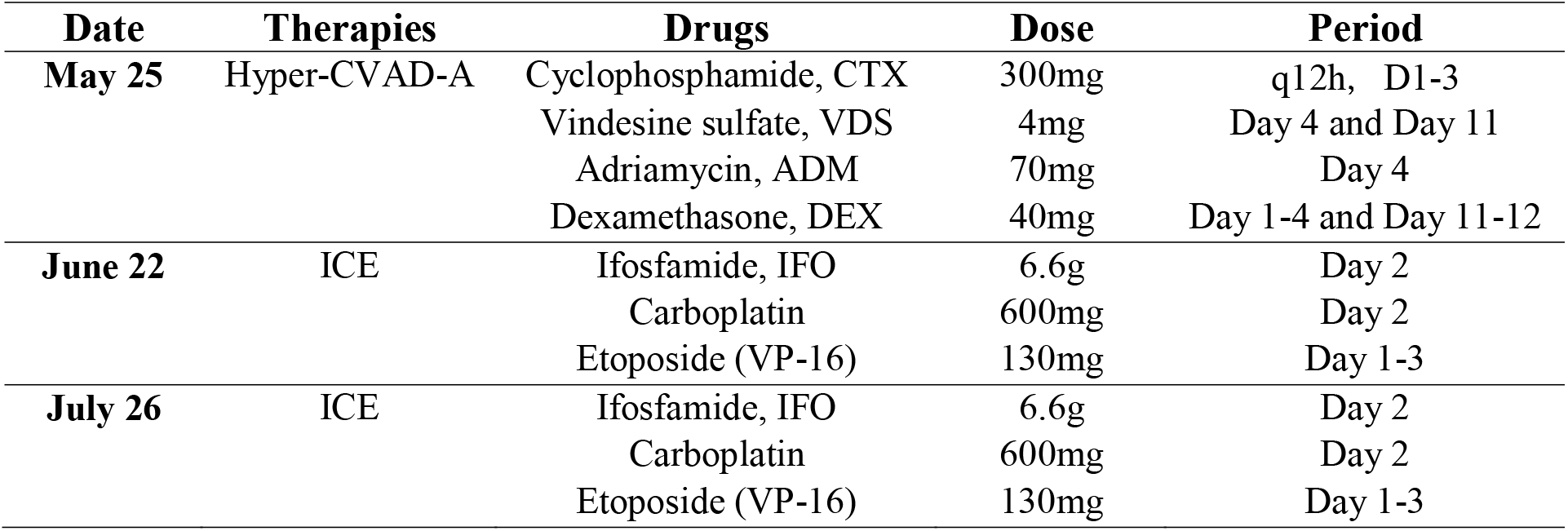
Chemotherapy administrations for the patient

| **Date** | **Therapies** | **Drugs** | **Dose** | **Period** |
| --- | --- | --- | --- | --- |
| **May 25** | Hyper-CVAD-A | Cyclophosphamide, CTX | 300mg | q12h, D1-3 |
|  |  | Vindesine sulfate, VDS | 4mg | Day 4 and Day 11 |
|  |  | Adriamycin, ADM | 70mg | Day 4 |
|  |  | Dexamethasone, DEX | 40mg | Day 1-4 and Day 11-12 |
| **June 22** | ICE | Ifosfamide, IFO | 6.6g | Day 2 |
|  |  | Carboplatin | 600mg | Day 2 |
|  |  | Etoposide (VP-16) | 130mg | Day 1-3 |
| **July 26** | ICE | Ifosfamide, IFO | 6.6g | Day 2 |
|  |  | Carboplatin | 600mg | Day 2 |
|  |  | Etoposide (VP-16) | 130mg | Day 1-3 |

### Establish a single cell atlas for HSTCL

To understand the cellular and molecular characteristics HSTCL and how they can be changed during chemotherapies, we performed paired scRNA-seq and scTCR-seq for 4 samples collected from the patient. 2 pre-treatment samples were collected from bone marrow (BM1) and PBMC (PBMC1), respectively, and 2 were collected post-treatment (BM2, PBMC2) (**Fig. 1A**). After quality control and data pre-processing, 36,092 cells (8,295 BM1, 9,189 PBMC1, 5,730 BM2 and 12,878 PBMC2) were retained for downstream analysis. Followed by dimension reduction and Leiden clustering, the cells were visualized by uniform manifold approximation (UMAP) (**Fig. 2A**). 7 major clusters were identified from the cell population, cell type identities were assigned to each cluster by canonical markers from previous publication (**Fig. 2B**): T and NK cells (T_NK, marked by CD3D, CD3E, NKG7 and GNLY), B cells (B, marked by CD79A and MS4A1), Plasma cells (Plasma, marked by MZB1 and IGKC), Monocytes (Monocyte1 and Monocyte2, marked by CD14, LYZ and CST3), Erythroid and progenitor cells (Ery_Pro, marked by HBD and CA2) and malignant γδ T cells (GD_Tumor, marked by TRCD, TRDV1 and CD3D). We didn’t observe strong batch effect among different samples (**Fig. S2**). To further confirm the identity of the tumor cluster, we evaluated the expression of HSTCL markers previously identified by immunohistochemistry and flow cytometry. We confirmed that these GD_Tumor cells were CD2+, CD3+, CD4-, CD5-, CD7+ and CD8, CD38, CD56 (NCAM1) partial positive, aligned with the diagnostic guidelines (**Fig. S3**)^4,6^.

**Figure 2.**
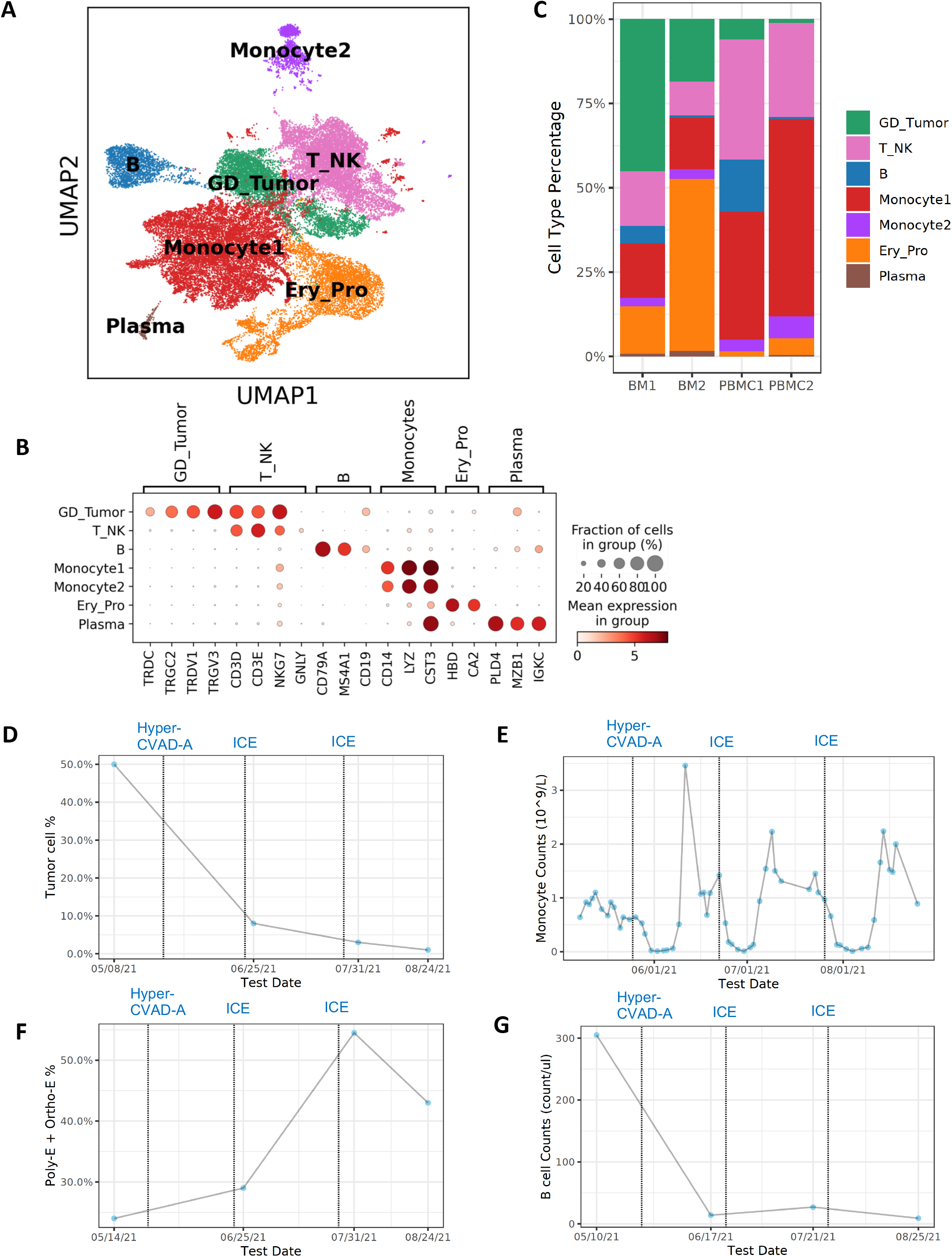
A cell type atlas of γδ hepatosplenic T-cell lymphoma. (A) Cell type atlas of γδ HSTCL presented by UMAP. (B) Dotplot of the canonical marker genes used to annotate major cell types. Dot size represents % of cells of that cluster expressing the given gene, while color indicates the expression level of that cluster. (C) Barplot to show the cell type compositions in each sequenced sample. Significant composition changes are observed pre- and post-treatment. (D-G) Cell type % or absolute count during treatments. Dotted lines indicate chemotherapy-initiated date. (D) Tumor cell % estimated from histology slides (Wright staining) at different time points. Tumor cell % is sharply reduced during the treatments. (E) Absolute monocyte counts flatulate during the disease progression, with a drop after chemotherapy administration and soon increase to an unusual high level. (F) In clinical bone marrow examinations, erythroid lineage cell % (polychromatophilic (Poly-E) and orthochromatophilic (Ortho-E)) shows an increasing trend. (G) In lymphocyte subtype counting, B cell absolute counts drop to very low level after treatment.

We observed unique cell type composition distribution in HSTCL and substantial composition changes post-treatment (**Fig. 2C**). Tumor cells were greatly reduced post-treatment in both PBMC (pre: 6.1% vs post: 1.1%) and BM (pre: 45.2% vs post: 18.5%), In Wright staining, tumor cell was also significantly decreased post-treatment (**Fig. 2D**). Unusual high percentage of monocytes was observed in PBMC samples, and became even higher post-treatment (pre: 40.2% and post: 65.0%, **Fig. 2C**), which was consistent with clinical blood examinations (**Fig. 2E**). Previous studies suggested that elevated monocyte count may associated with poor prognosis and identify high-risk patients in lymphomas^15,16^. Ery_pro cell % were increased post-treatment in both BM (pre:14.2% vs post: 51.1%) and PBMC (pre:1.6% vs post: 5.0%). In Ery_pro cluster, most cells were erythroid and erythroid progenitor cells (**Fig. S4**). In clinical bone marrow examination, erythroid lineage cell % also showed an increasing trend (**Fig. 2F**). B cells reduced to a very low percentage post-treatment in both BM (pre: 5.2% vs post: 0.8%) and PBMC (pre: 15.7% vs post: 0.7%). In lymphocyte subtype counting by flow cytometry, B cells were significantly dropped after chemotherapy (**Fig. 2G**). These results suggested that the tumor microenvironment (TME) was dynamically reshaping as responses to disease progression and treatment.

### HSTCL malignant γδ T cells had unique clonal and molecular features

To investigate the clonal expansion of the malignant γδ T cells, we performed scTCR-seq in parallel with scRNA-seq. Among the 5,515 cells in the GD_Tumor cluster, 4,505 (81.7%) cells had recoverable γδ TCR information. The TCR repertoire of malignant cells was almost founded by a single clonotype, suggesting the tumor cells likely arose from a single ancestor cell and clonal expanded subsequently (**Fig. 3A**). For the 4,419 cells had assembled γ chains, 4,374 (99.0%) had the same γ chain sequence (TRGV3 + TRGJ1 + CATWDNLMENYYKKLF), for the 3,614 cells had assembled δ chains, 3,576 (99.0%) had the same δ chain sequences (TRDV1 + TRDJ1 + CALAVVVGGLGTDKLIF), and for the 3,494 cells had both γ chain and δ chain assembled, 3614 cells had the above γ chain and δ chains sequences (**Fig. 3A** blue dots). As a comparison, we identified 559 cells in NK_T cluster with γδ TCR, suggesting they were normal γδ T cells from this patient. These normal γδ T cells had a more diverse TCR repertoire, ~60% of the cells were singletons (**Fig. 3B**).

**Figure 3.**
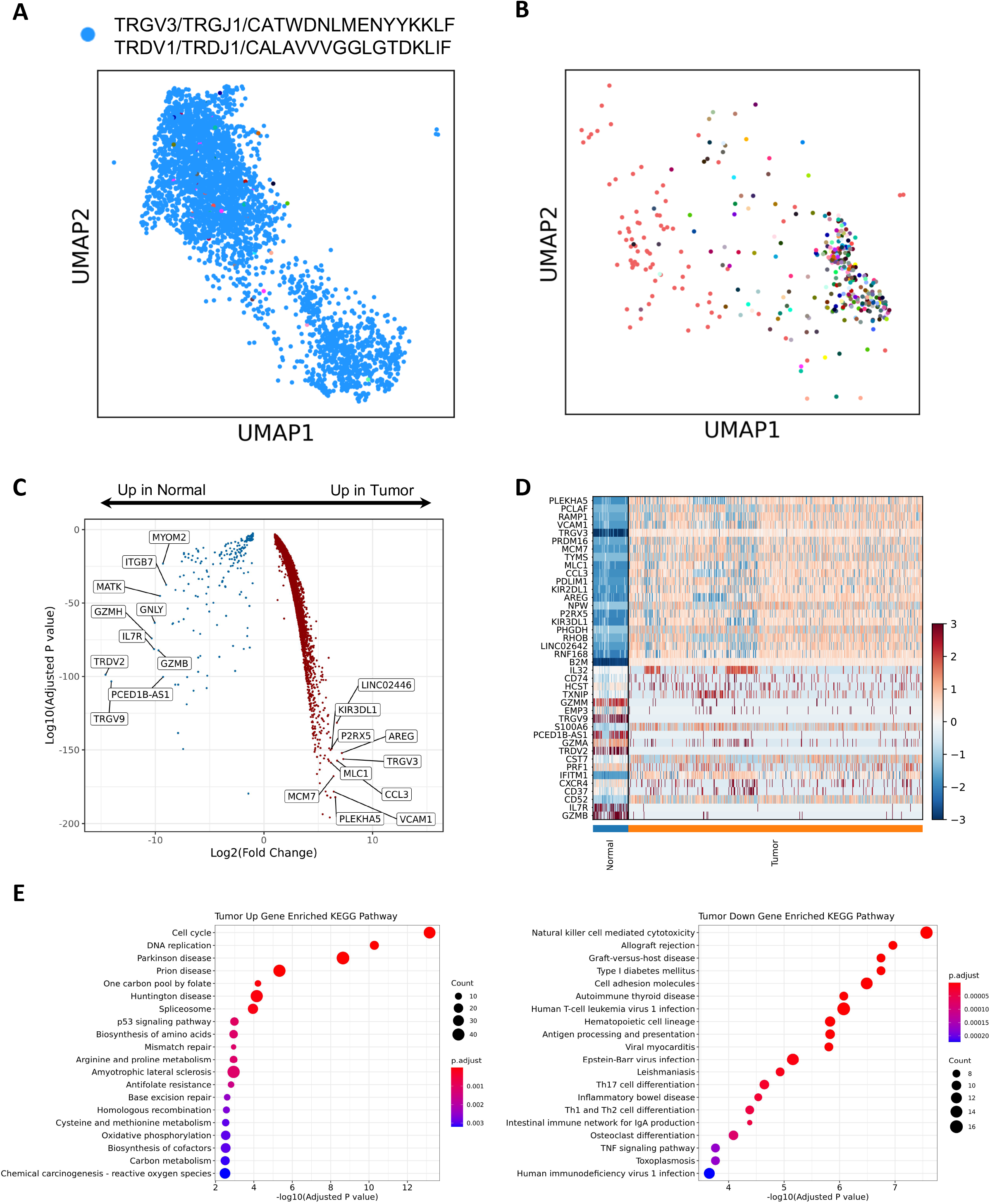
Molecular features of γδ hepatosplenic T-cell lymphoma. (A) TCR sequencing reveals the distribution of a single prominent clonotype in the malignant γδ T cells (Blue dots). (B) TCR sequencing shows that the normal γδ T cells have a diverse TCR repertoire, colors indicate different clonotypes. (C) Volcano plot to show the differentially expressed genes between malignant and normal γδ T cells, top 10 up and down genes by fold changes are labeled. (D) Heatmap to present the top 20 differentially expressed genes between malignant and normal γδ T cells. (E) KEGG pathway enrichment of the differentially expressed genes. Dot size represents the number of significantly changed genes in the pathway and color indicates adjusted p values.

To give a more comprehensive description of the malignant γδ T cells and search for novel candidate genes relevant to HSTCL pathogenesis, we compared the malignant γδ T cells gene expressions with those of normal γδ T cells. We found a substantial number of genes were differentially expressed between the malignant and normal γδ T cells (**Fig. 3C, Table. S1**). Among them, the top up-regulated genes in tumor cells were those related to cancer progression (eg, AREG, CCL3, PLEKHA5, VCAM1 and MCM7, **Fig. 3D**), which might be the targets of innovative treatments. Normal γδ T cells have strong cytotoxicity by expressing NK cell-associated genes like NKG2D, GNLY and a profile of cytotoxic molecules (eg, GZMA, GZMB, GZMM, and PRF1)^17–19^. However, these genes were mostly absent in the malignant γδ T cells (**Fig. S5**). On the contrary, killer cell immunoglobulin-like receptor 2DL2 (KIR2DL2) and KIR2DL3 were increased in the malignant γδ T cells, which encode the inhibitory receptors of the KIR family and reduce the cells’ cytotoxic function^20^. These results suggested that malignant γδ T cells differed from normal γδ T cells by lacking the cytotoxic modules. KEGG pathway enrichment analysis further showed that the top 1000 tumor up-regulated genes were significantly enrichment in pathways involving in cell proliferation and cancer development, such as Cell cycle, DNA replication, p53 signaling pathways (Adjusted p value = 6.99e-14, 5.28e-11 and 1.03e-03, respectively. **Fig. 4E**).

**Figure 4.**
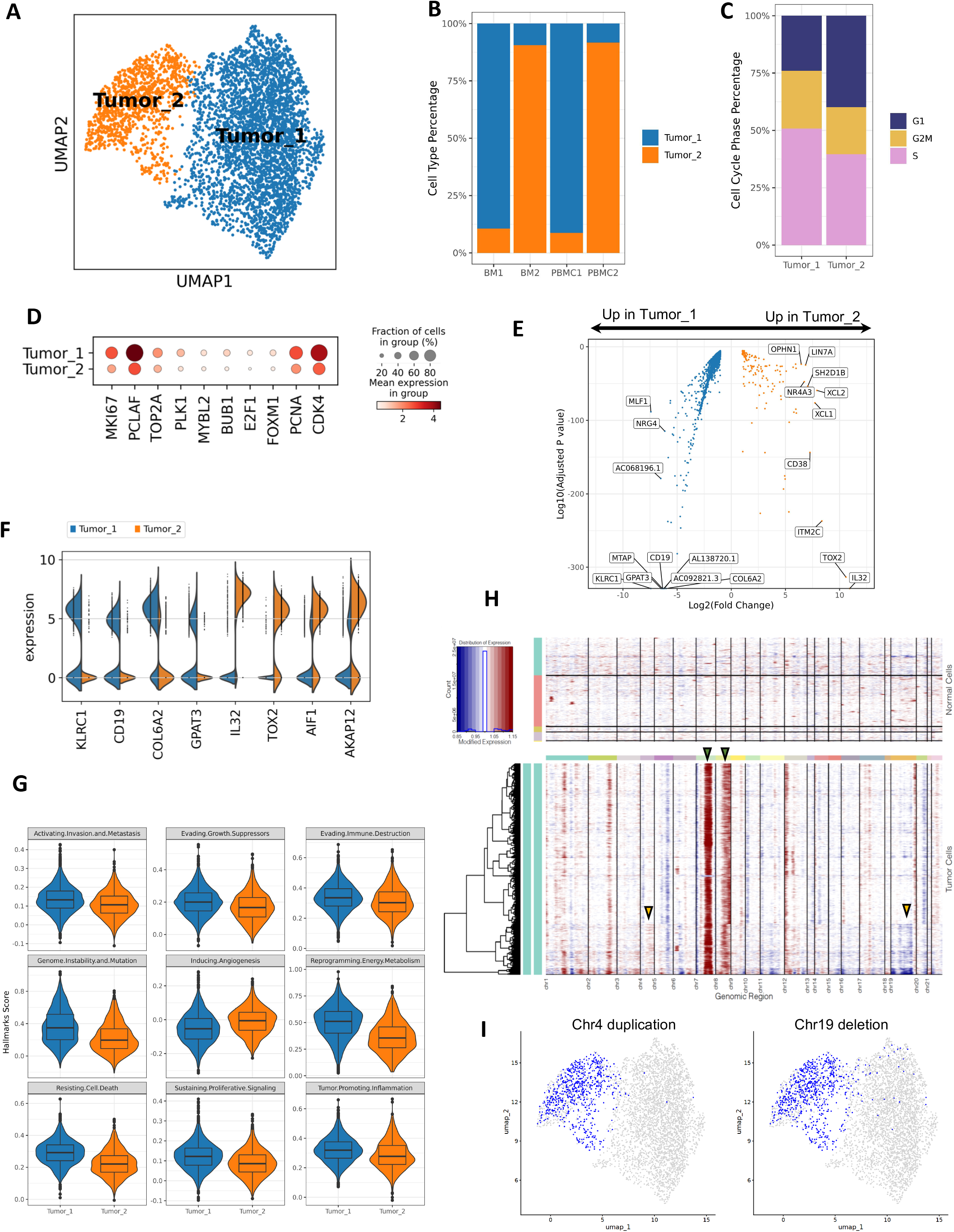
scRNA-seq reveals tumor cell heterogeneities. (A) UMAP of malignant γδ T cells, which are separated into two distinct sub-clusters. (B) Barplot to show that Tumor_1 cells are mostly from pre-treatment samples (BM1 and PBMC1) while Tumor_2 cells are dominant in post-treatment samples (BM2 and PBMC2). (C) Cell cycle phase distribution among Tumor_1 and Tumor_2. Tumor_1 has a higher portion of cells in S and G2M phases. (D) Dotplot of proliferation markers expression in Tumor_1 and Tumor_2. (E) Volcano plot to show the differentially expressed genes between Tumor_2 and Tumor_1 cells, top 10 up and down genes by fold changes are labeled. (F) Violin plot of selected differentially expressed genes between Tumor_1 and Tumor_2. (G) Violin and box plot of Cancer Hallmarks scores. Tumor_1 has higher scores in most cancer hallmarks. (H) Heatmap of the inferred CNVs in Tumor cells using normal immune cells as references. Red indicates CNV gains and blue indicates CNV loss. Green arrows indicate chromosome 7 and 8 duplications inferred in majority of the tumor cells. Yellow arrows indicate tumor sub-cluster specific CNVs. (I) Selected tumor sub-cluster specific CNVs. Left: a chromosome 4 partial duplication is enriched in Tumor_2. Right: a chromosome 19 partial deletion is enriched in Tumor_2. Blue indicates cells with specified CNVs.

### Transcriptional landscape revealed the heterogeneity of HSTCL malignant cells

We next investigated the heterogeneity of the malignant cells at single-cell level. The 4505 cells from GD_Tumor cluster with γδ TCR were taken out and re-clustered. Two distinct tumor clusters were identified from these cells (**Fig. 4A**), which had a strong correlation with the treatment status (**Fig. 4B**). Tumor_1 cluster consisted 3439 (75.3%) tumor cells, 98.0% of which were from pre-treatment samples (BM1 and PBMC1) and were almost absent in post-treatment samples. Tumor_2 cluster was relative smaller, included 1066 cells but dominating the post-treatment samples (BM2 and PBMC2), suggesting the chemotherapies may have a selection effect on the tumor cell populations.

We further did a comprehensive comparison between the two tumor clusters. From the expression of G2M and S-phase genes, we inferred the cell cycle phases for each tumor cell and observed different phase distribution between the two clusters (**Fig. 4C, Fig. S6**). 76.0% of the Tumor_1 cells were at S and G2M phases, while only 60.1% in the Tumor_2 cluster. We then surveyed the expression of proliferation associated genes (eg, MKI67, PCLAF, PCNA), Tumor_1 cells also had higher expression than Tumor_2 among these genes (**Fig. 4D**), together with the cell cycle analysis, suggesting that Tumor_1 cells were more actively proliferating than Tumor_2 cells.

Differentially expressed genes were identified from the two clusters (**Fig. 4E** and **F**, **Table. S2**). NK cell receptors (NKR) were frequently expressed in normal γδ T cells. We found the kill receptor KLRC1 (NKG2A) was expressed in 62.0% Tumor_1 cells but only 3.3% in Tumor_2. KLRK1 (NKG2D), another NKR highly expressed in normal γδ T cells, was almost absent in Tumor_1 but presented in 25.8% Tumor_2 cells. Unexpectedly, we observed high expression level of CD19, a marker for B cell, in Tumor_1. Other B cell marker such as CD79, CD20 were not identified in Tumor_1 (**Fig. S7**), suggesting the CD19 expression was not caused by doublet. Therefore, Tumor_1 had a CD3, CD19 double positive phenotype, which is rarely reported in previous studies^21,22^. In Tumor_2, the tumor cells lost the CD19 expression. Among the genes highly expressed in Tumor_2, many were associated with tumor progression and potential drug resistance. IL32 was reported to accelerates the proliferation of cutaneous T cell lymphoma cell lines through mitogen-activated protein kinase (MAPK) and NF-κB signaling pathways^23^. TOX2 is a novel tumor driver, which promotes in Natural Killer/T-Cell Lymphoma cell growth and enhances ability of colony formation, as well as protects cell viability under adverse condition^24^. AIF1 and AKAP12 were reported to correlated with various chemo resistance^25,26^. These results suggested that Tumor_2 might obtained unique survival features under the therapeutic conditions.

To describe different phenotypic characteristics of the malignant cells, we calculated the 10 cancer hallmarks scores curated by Cancer Hallmark Genes (CHG) database^27^. By permutation test, we identified 9 out of the 10 hallmark scores were significantly different between the two tumor clusters (**Fig. 4G, Table. S3**). Except “Inducing Angiogenesis”, all other 8 hallmarks were significantly higher in Tumor_1 cluster, indicating Tumor_1 had stronger cancer related signaling pathways. Since Tumor_2 cells were mostly from post-treatment samples, the decreased hallmarks scores could be a response to therapeutic pressure. Tumor_2 might be a less aggressive tumor subtype, however, may have unique features to tolerant the chemotherapy.

Chromosomal aberrations are frequently observed in HSTCL^28,29^. To determine if the transcriptional heterogeneity was potentially associated with chromosomal abnormalities, we inferred large scale copy number variations (CNV) at single cell resolution using inferCNV R package. We identified gains of chromosome 7 and 8 in majority of the tumor cells (**Fig. 4H** green arrows), which are the most common chromosomal abnormalities in HSTCL, presented in up to 63% and 50% of cases, respectively, and frequently cooccurring^28,29^. The two tumor clusters also showed cluster-specific CNVs. For example, a portion of cells in Tumor_2 cluster demonstrated a CNV gain in chromosome 4 and a loss in chromosome 19 (**Fig. 4H** yellow arrows and **Fig. 4I**). These findings suggested that some chromosomal aberrations might have occurred during the evolution of the tumor cells under therapeutic pressures and may contribute to the survival of the tumor cells.

### Tumor microenvironment heterogeneity associated with disease progression

To characterize the features of TME in this HSTCL patient, we performed analysis for the immune cells in the single cell profiles, with a focus on T cells and monocytes. We first re-clustered the cells in T_NK cluster, and grouped them into 4 major sub-clusters. Based on the expression signatures, these clusters can be annotated as CD4 T cells (CD4_T), CD8 memory T cells (CD8_Mem_T), CD8 naïve T cells (CD8_Naive_T) and NK cells (NK) (**Fig. 5A** and **B**, **Fig. S8**). In the post-treatment samples, the CD8 memory T cell % were increased (BM1 21.9% vs BM2 40.6% and PBMC1 21.7% vs PBMC2 35.3%, **Fig. 5C**), indicating these cells populated during disease progression. scTCR-seq results showed that the cells in CD4_T and CD8_Naive_T were mostly singletons (71.5% and 73.0%, respectively). Meanwhile, CD8_Mem_T cluster had more clonal expanded cells, only 40.6% were singletons, consisting with their cytotoxic and memory features (**Fig. 5D**). Cells in the post-treatment samples were more clonal expanded than the pre-treatment samples (**Fig. 5E**). In addition, we observed multiple clonotypes were already detected in pre-treatment samples then further expanded in the post-treatment samples (**Fig. 5F**), suggesting they were potential effector cells, expanding during the disease progression to perform anti-tumor functions.

**Figure 5.**
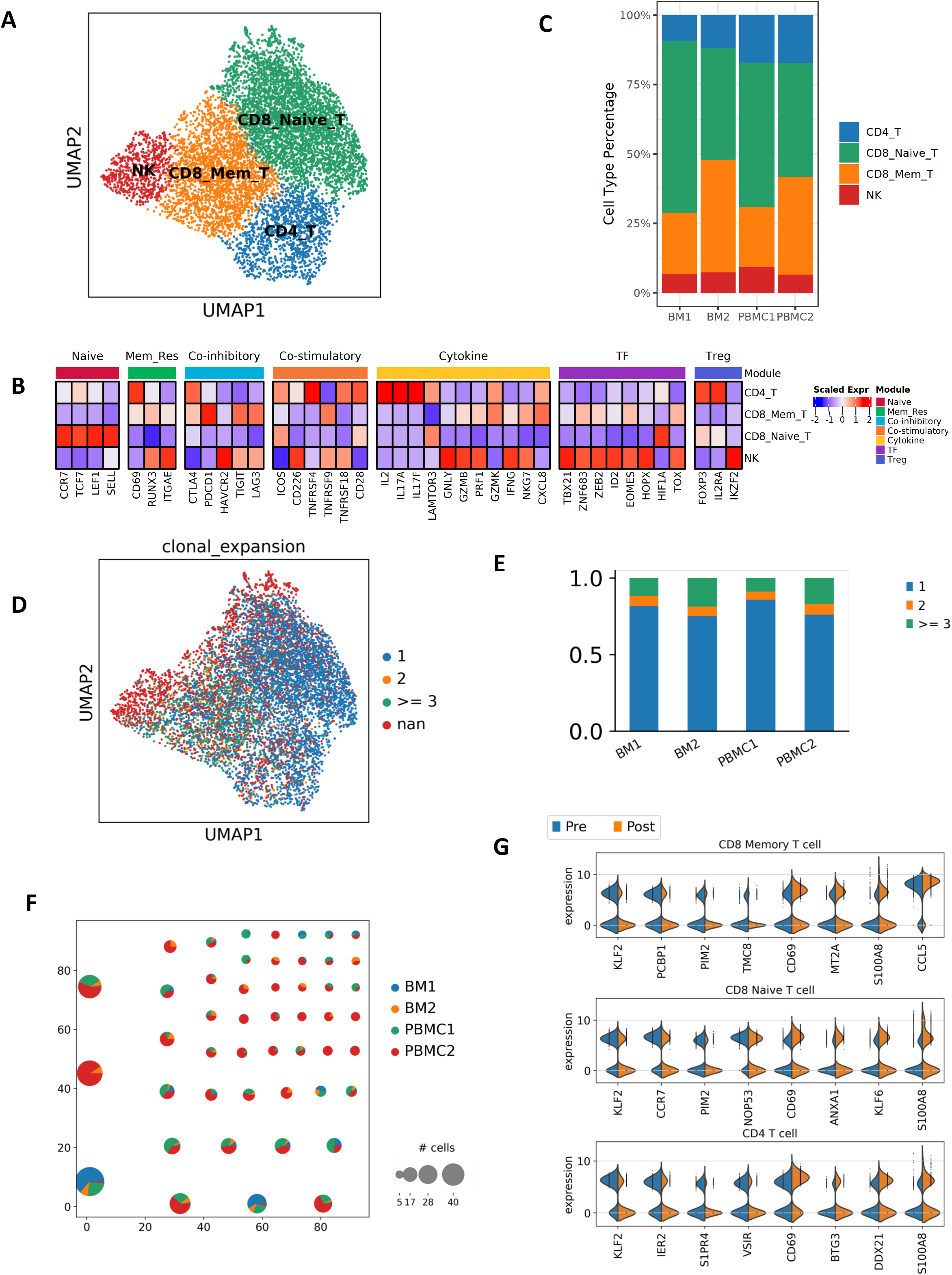
T cell subtypes and dynamics in tumor microenvironment. (A and B) T cell subtypes are identified by canonical markers and unique cell type features. (C) T cell subtype composition changes during the disease progression. CD8 memory T cells are increased post-treatment. (D) Clonal size distribution among the T cells. Clonal expanded cells are enriched in CD8 memory T cells. (E) Barplot to show the clonal size distribution in each sequenced sample, post-treatment samples have less singletons, suggesting lower diversity (F) Pie plot to show the clonotypes with clonal size > = 5. Pie size represents the size of the clonotype, color indicate the cell origins. (G) Selected differentially expressed genes between post- and pre-treatment cells in each T cell subtype.

We next analyzed differentially expressed genes in T-cell subsets pre- and post-treatment (**Fig. 5G, Table. S4-6**). We observed noticeable changes in the transcriptomes of these cells, involving genes that related to tumor invasion, metastasis, and therapeutic resistance. For example, CD69 was increased post-treatment in all three T cell subsets. CD69 was demonstrated to regulate T-cell migration and retention in tissues, playing an important role in inducing the exhaustion of tumor-infiltrating T cells^30^. ANXA1 was increased in CD8 memory T cell post-treatment, which has been reported to inhibit the anti-tumor immunity and support the formation of an immunosuppressed tumor microenvironment that promotes tumor growth and metastasis^31^.

We observed substantial increase of monocytes in the PBMC samples (**Fig. 2C**), suggesting monocytes may also play a role in the disease progression. Here, we further evaluated the heterogeneity of the monocyte population. We identified two major monocyte subtypes, the classic monocytes (Classic_Mono) and the CD16+ Monocytes (CD16_Mono) (**Fig. 6A** and **B**). The CD16+ nonclassical monocytes might have a function to scavenge tumor cells and debris due to their patrolling activities^32^. It accounted 25.4% and 58.5% of the monocyte population in BM1 and PBMC1, respectively, but decreased to 9.9% and 26.3% post-treatment. These findings demonstrated the actively reforming TME that may influence tumor development and therapeutic outcomes.

**Figure 6.**
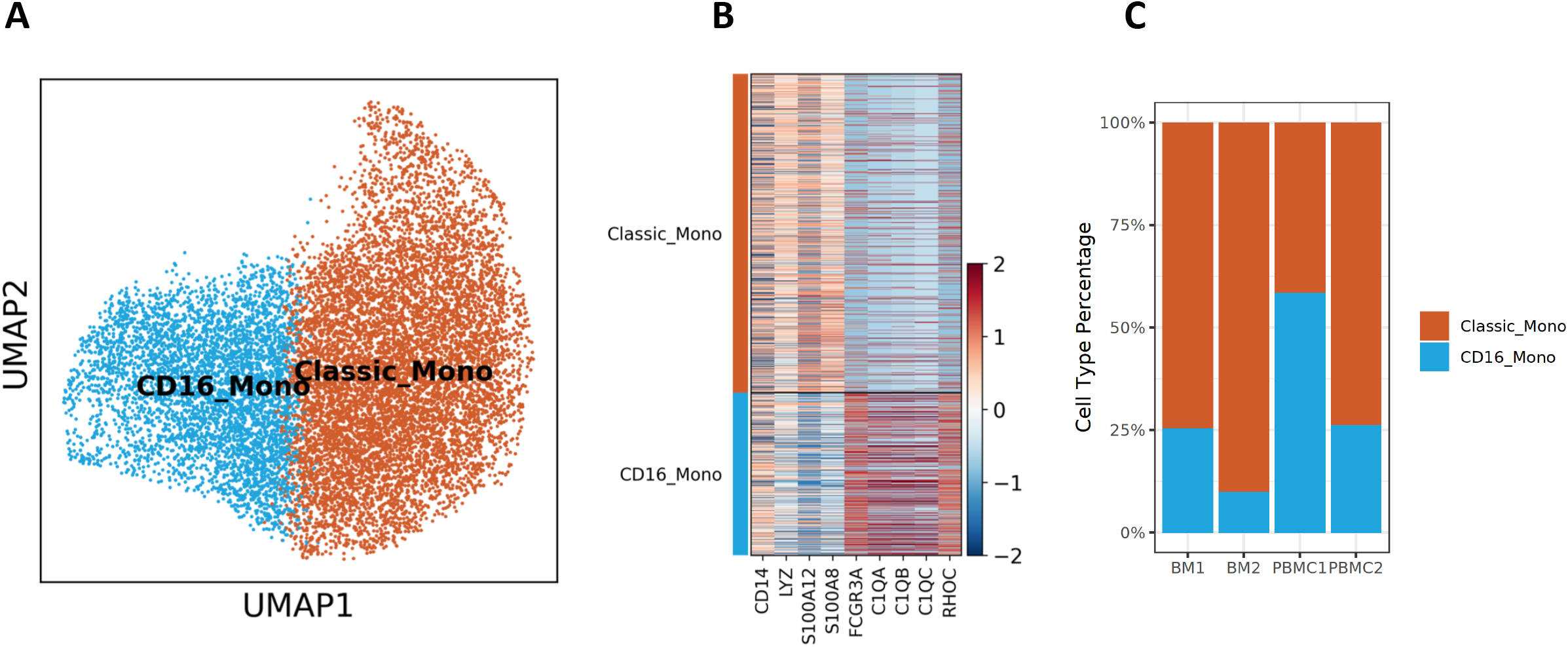
Monocyte subtypes and composition changes in tumor microenvironment. (A) UMAP presentation of monocyte subtypes. (B) Heatmap to present the canonical marker genes used to identify the classical monocytes and the CD16+ nonclassical monocytes. (C) Barplot to show the monocyte subtype composition changes pre- and post-treatment. CD16+ nonclassical monocytes decreased post-treatment.

### Rewired Tumor-microenvironment interaction networks during treatment

As we observed substantial changes in TME during the disease progression, we further performed cell-cell interaction analysis to explore the communication dynamics between tumor cells and other TME cells. We revealed significantly decreased cell-cell interactions in post-treatment cells, for both total interaction counts and interaction strengths (**Fig. 7A** and **B**). The total interaction counts were 1487 in BM1 cells, decreased to 913 in BM2 cells. Among them, tumor cell involving interactions decreased from 709 to 418, contributing to 50.7% of the decrease (**Fig. S9**). We also observed that several signaling pathways were affected by the interaction network rewiring (**Fig. 7C**). For example, cell-cell interactions in MHC-1, MIF, GALECTIN and ICAM signaling pathways became weaker in post-treatment cells. We further evaluated each ligand-receptor pair between tumor and other major immune cells (**Fig. 7D**). As examples, MIF-(CD74+CXCR4) interaction was decreased between tumor and most immune cell types post-treatment. MIF/CXCR4 axis were reported to induce an aggressive phenotype by inducing proliferation, adhesion, migration, and invasion of the colon cancer cells^33^. ADGRE5 (CD97)-CD55 interaction were also decreased post-treatment, an interaction has been shown to contributing to aggressiveness in multiple cancers^34–36^. Together, our results suggested that the interaction between malignant γδ T cells and the TME were actively reshaping during the disease progression. The Tumor_2 cells had cut down the communications with the TME, however might acquire survival advantages.

**Figure 7.**
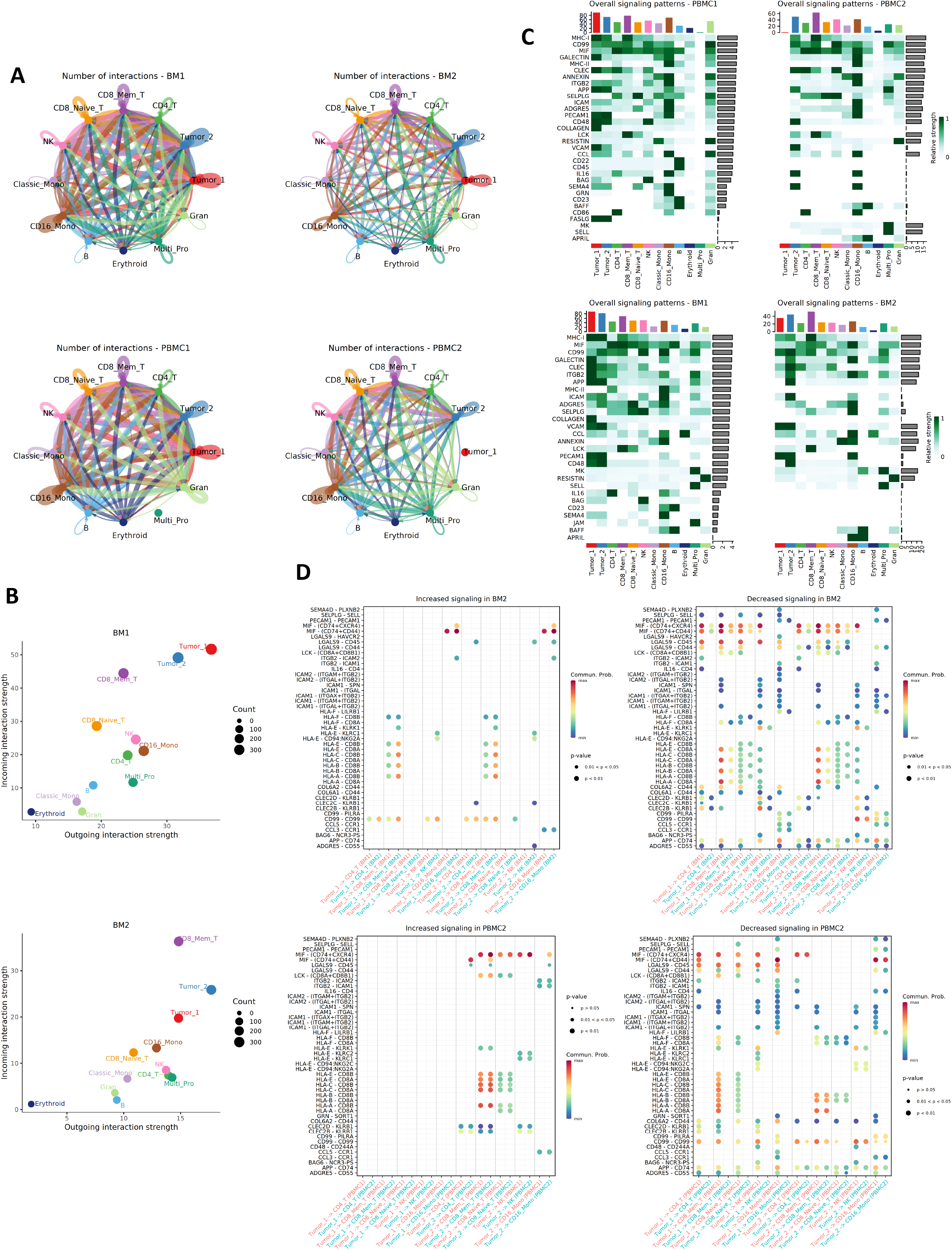
Rewired cell-cell interaction networks during disease progression. (A) Overview of the cell-cell interactions inferred from ligand-receptor expressions. Post-treatment samples have less interactions than the pre-treatment samples. (B) Scatter plot of the incoming and outgoing signals for each cell type. Tumor cell signals are reduced post-treatment. (C) Heatmap to compare the signaling patterns pre- and post-treatment. (D) Bubble plot to show the increasing and decreasing ligand-receptor interactions between tumor cells and major immune cells. Bubble sizes indicate p values and colors indicate average expression of ligand and receptor in the given interaction.

## Discussion

HSTCL is a highly aggressive tumor with limited knowledge of its pathogenesis. In this study, we provided the first single cell transcriptome and T cell receptor atlas of HSTCL at both treatment naïve status and post-chemotherapy. Although limited to a single patient, our study provides a framework of using single-cell technologies to dissect disease in a clinically meaningful way, providing plentiful information for future diagnosis and personalized treatment of this disease. By comparing with normal γδ T cells, we described the molecular signatures of HSTCL. We found that the malignant γδ T cells didn’t resemble the cytotoxic modules as the normal γδ T cells. Cytotoxic molecules like granzyme and perforins were negative or only lowly expressed. On the contrary, several oncogenesis-related gene were highly expressed in the malignant γδ T cells: AREG, of which the oncogenic activities were well documented in many cancer types, including hematological cancers^37^. PLEKHA5 has been identified to regulates tumor growth in metastatic melanoma and gastric carcinoma^38,39^. Our findings suggested these gene may also involve in HSTCL pathogenesis and offer potential novel therapeutic targets.

Single cell technology also allowed us to evaluate the heterogeneity of the HSTCL tumor cells. Although the TCR sequencing data suggested that 99% of the tumor cells carried the same TCR thus should be derived from the same ancestor, we found the tumor cells evolved into two transcriptionally distant subtypes. The pre-treatment tumor cells were dominated by Tumor_1, which presented a more aggressive molecular phenotypes than Tumor_2 cells. Though the overall tumor cell load decreased post-treatment, Tumor_2 cells were expanded and became the dominant tumor subtype, while Tumor_1 cells were almost disappeared. Differential gene expression analysis identified noticeable differences between the two tumor subtypes, with multiple genes related to lymphoma progression and chemo-resistance were significantly increased in Tumor_2. IL32 exerts both anti-tumor or pro-tumor effects in a cancer type dependent manner^40^ and it has been reported to play a role in the formation and maintenance of lesions in cutaneous T-cell lymphoma (CTCL)^23^. TOX2 was recently reported to be a novel tumor driver in natural killer/T-cell lymphoma (NKTL)^24^ and high expression of TOX2 was associated with worse overall survival in both NKTL and acute myeloid leukemia (AML)^24,41^. AIF1 and AKAP12 were associated with chemo-resistance. AIF1 has been demonstrated to reinforce the resistance of breast cancer cells to cisplatin by inhibition of cell apoptosis and reduction of intracellular cisplatin accumulation^26^ and AKAP12 is associated with paclitaxel-resistance in serous ovarian cancer^25^. Together, these results suggested that Tumor_1 cell was highly aggressive but might be more sensitive to the administrated chemotherapies while Tumor_2 cells could harbor unique features to tolerant the therapeutic pressures. Interestingly, we also observed high expression of CD19 in Tumor_1 but not in Tumor_2 cells. CD19 co-expression in a mature T cell neoplasm has barely been reported before. Rizzo et al. reported the first case of CD19 expression in peripheral T-cell lymphomas (PTCL)^22^ and Jain et al reported a rare case of CD19 positive HSTCL^21^.Our observations suggested the CD19 positive and negative T cell neoplasm can co-exist within the same patient and potentially related to the drug sensitivity. In addition to gene expression profiles, we also identified subtype-specific CNVs among the tumor cells, adding another layer of heterogeneity of the tumor population. Currently, there is no efficient treatment for HSTCL. The heterogeneity, specifically the arising of multiple malignant subtypes, may improve our understanding of the disease refractory, where malignant subtypes can be resistant to treatment.

Recent studies have highlighted the critical role of TME in the evolution of various tumors^42,43^. In this study, we also observed dynamic reforming of the TME during the disease progression. We noticed significant increasing of monocytes percentage among the immune cells, and further dissected the heterogenicity of the monocyte population. During cancer, different monocyte subsets can perform both pro- and antitumoral function^32^. Our results indicated that the composition of monocyte subtype was rebalanced during the disease. The T cell population was also reshaped pre- and post-treatment, in both TCR repertoire and gene expression profiles. Post-treatment T cells had more clonal expanded cells that may have interacted with tumor cells. On the other side, the reformatted T cell transcriptomes may also promote evolution of malignant subtypes.

In summary, our study generated a comprehensive view of molecular heterogeneity and tumor microenvironment in HSTCL, provided a valuable resource for understanding the pathogenesis of HSTCL and exploring the potential therapeutic targets in the future.

## Acknowledgments

We thank Mr. Wei Wang and Drs. Tao Yang, Jing He and Michelle Dow for their insightful discussions and comments on the manuscript. We thank the Hematology Diagnostic Laboratory in the 1st People’s Hospital of Yunnan Province for their supports of clinical examinations. We thank Sinotech Genomics Co., Ltd. (Shanghai, China) for their support for single cell experiments. This study was supported by research funding from Yunnan Province Clinical Center for Hematologic Disease. Haixi Zhang was supported by grant from Yunnan Province Clinical Center for Hematologic Disease (2020LCZXKF-XY16).

## Authorship Contributions

RZ, KS and WS initiated the project and designed the experiment with input from HZ. WS and HZ performed the majority of experiments. HZ, KS involved in patient recruitment and sample collection. WS analyzed the CT scanning data. RZ performed the bioinformatics work. RZ, KS, WS, HZ, FY, KN and CW interpreted the data and results. RZ wrote the initial draft with all authors providing input on the manuscript.

## Conflict of Interest Disclosures

RZ is a paid consultant of Innovec Biotherapeutics. CW is the co-founder of Innovec Biotherapeutics.

