## Supplement Figures for "Single cell profiling of γδ hepatosplenic T-cell lymphoma unravels tumor cell heterogeneity associated with disease progression"

**Figure S1. CT scans of lymph node indicate lymphadenopathy is absent.** (A) Red marks left inguinal lymph node (B) Red marks left upper cervical lymph node

A

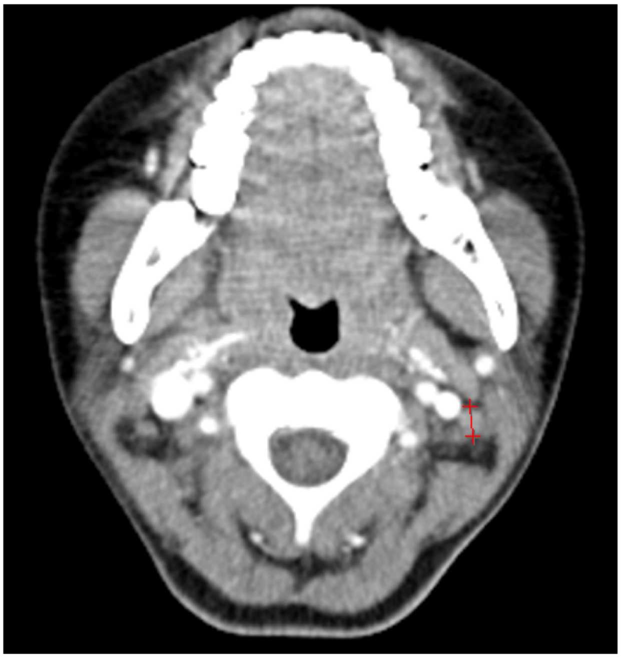

B

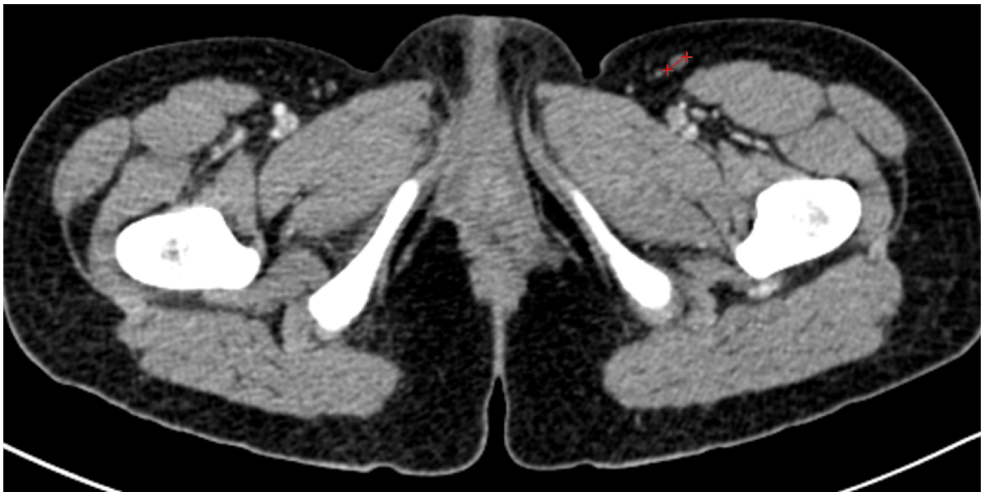

**Figure S2. UMAP projections of each sequenced sample.**

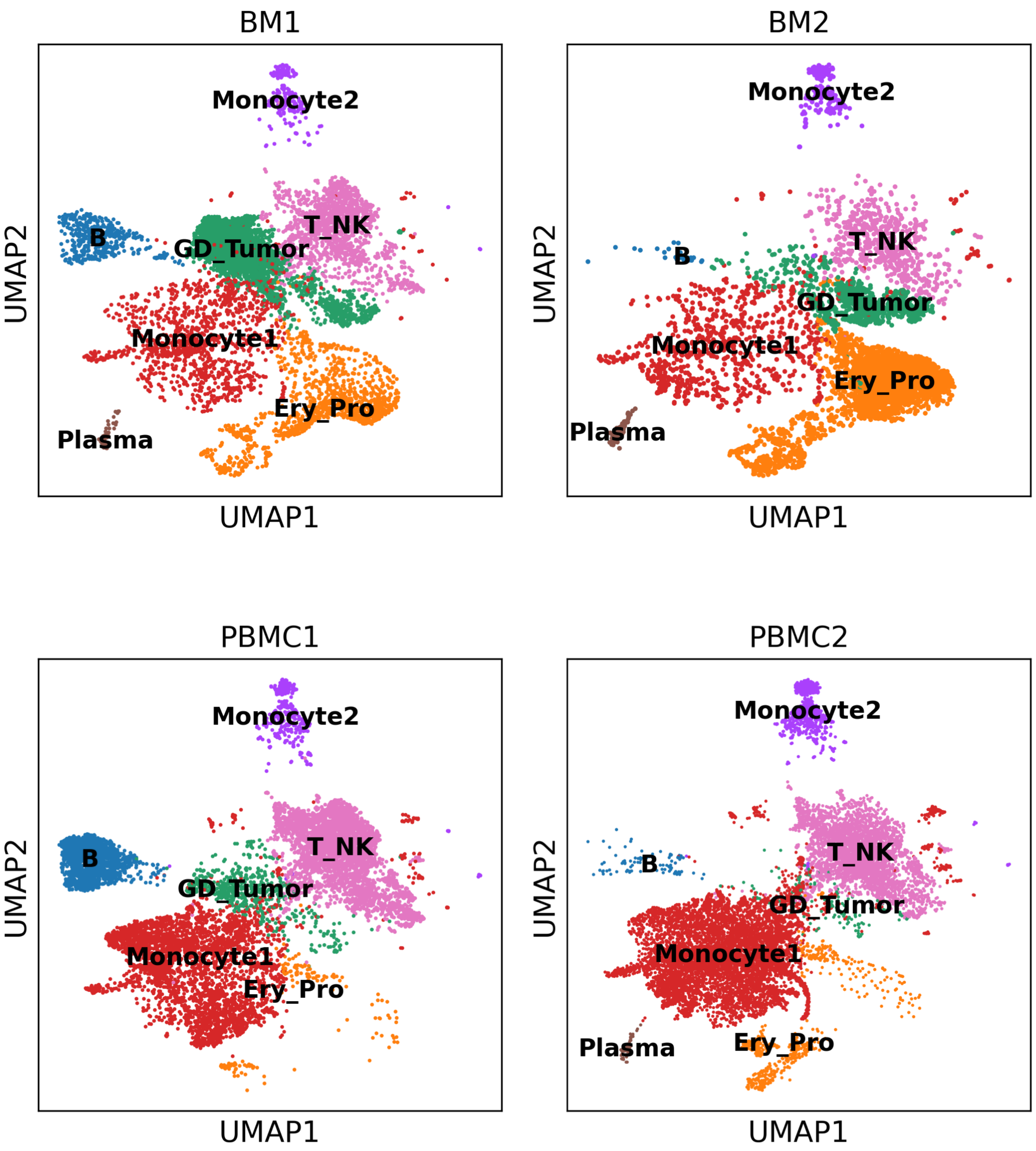

### Figure S3. Expression of common markers used for HSTCL diagnosis

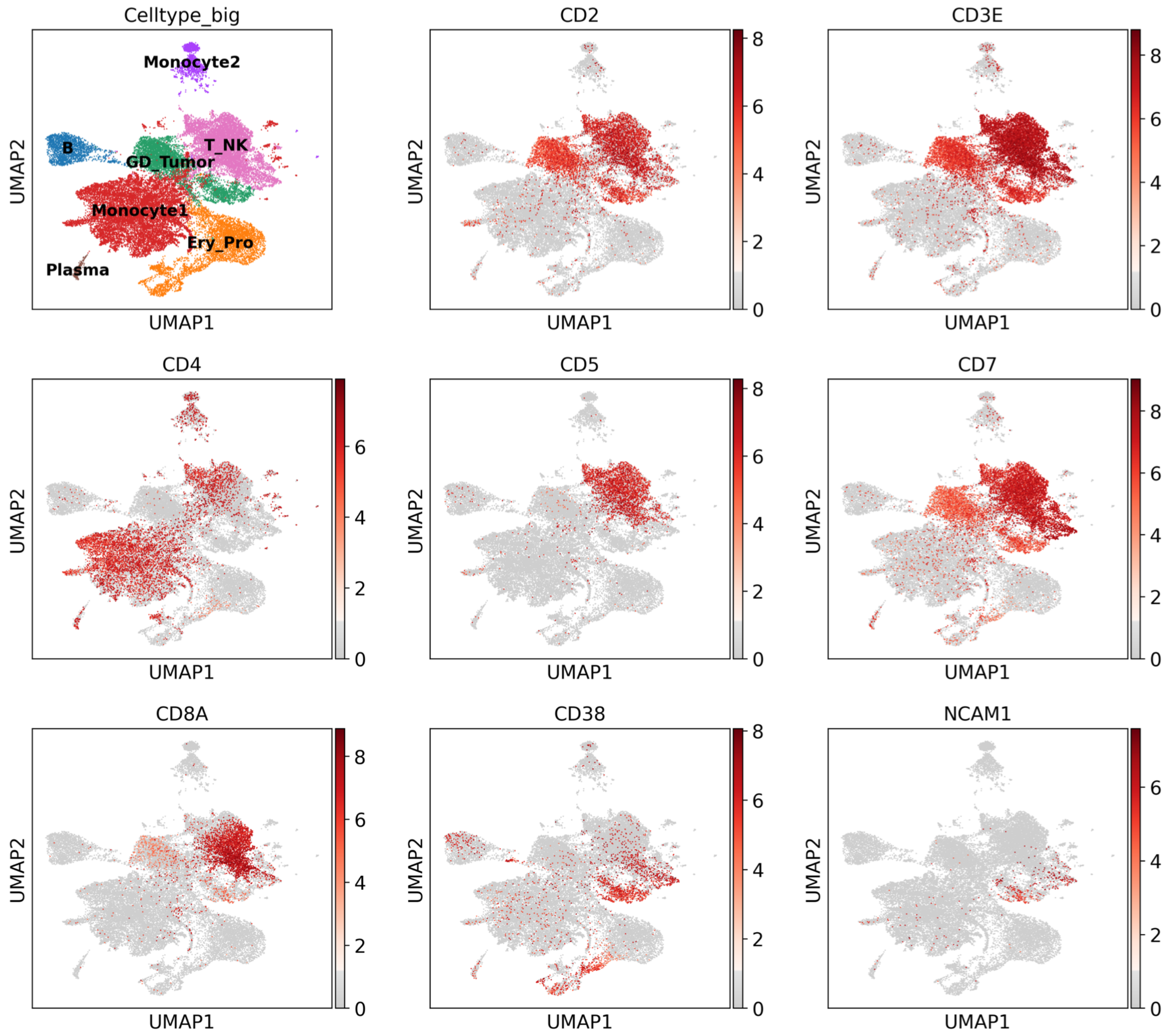

**Figure S4. Sub-clusters identified in Ery_Pro.** (A) UMAP of sub-clusters in Ery_Pro. Gran: Granulocyte progenitors, Multi_Pro: multipotent progenitors (B) The expression of marker genes for to the sub-clusters annotation

A

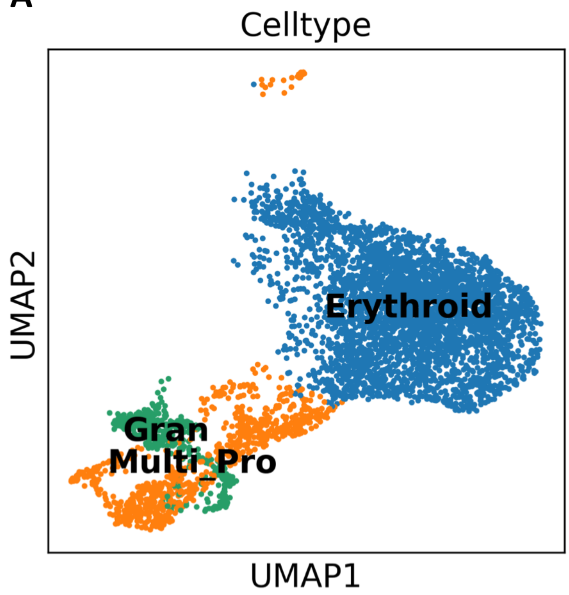


B

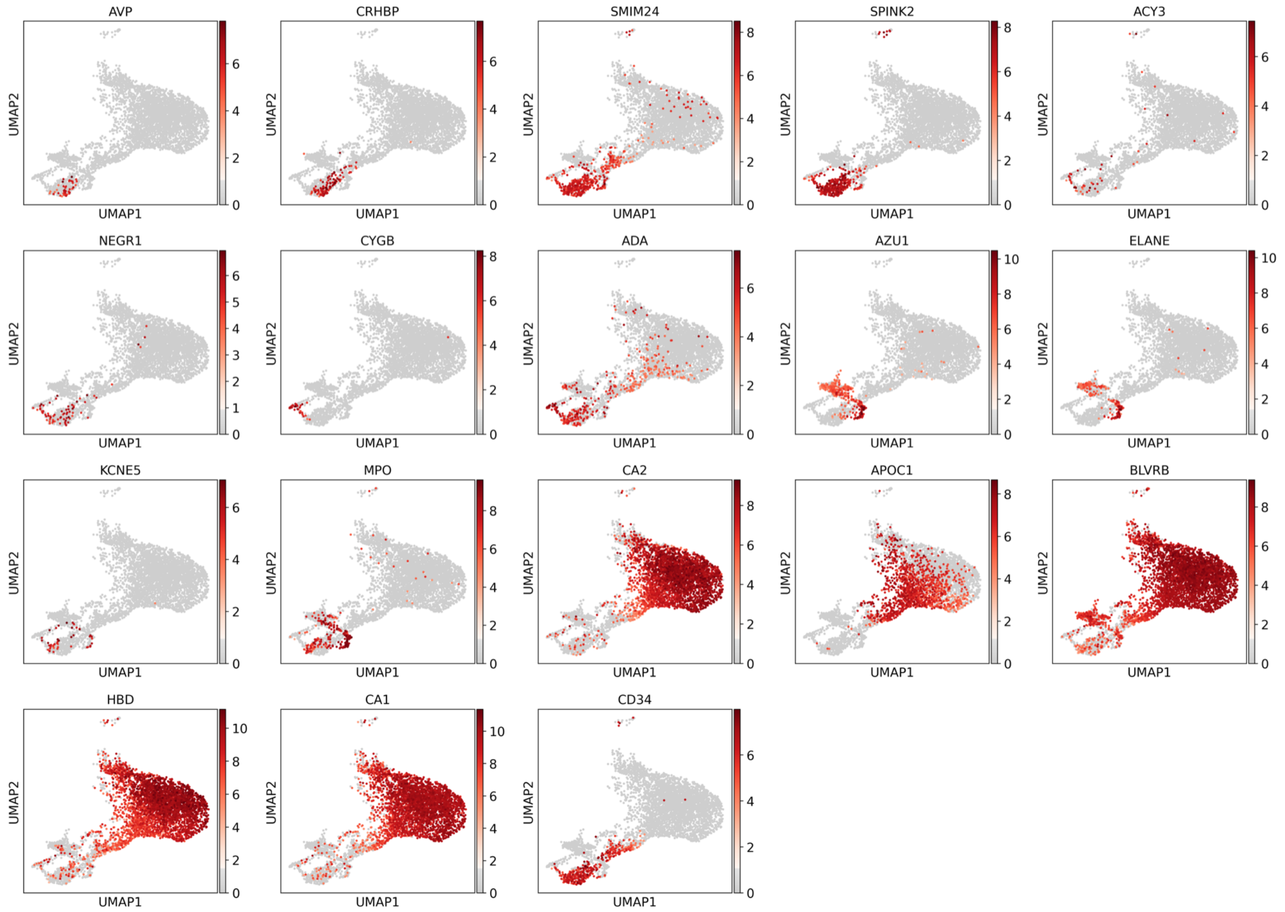

### Figure S5. Cytotoxic genes reported to be expressed in normal γδ T cells had no or low expression in malignant γδ T cells

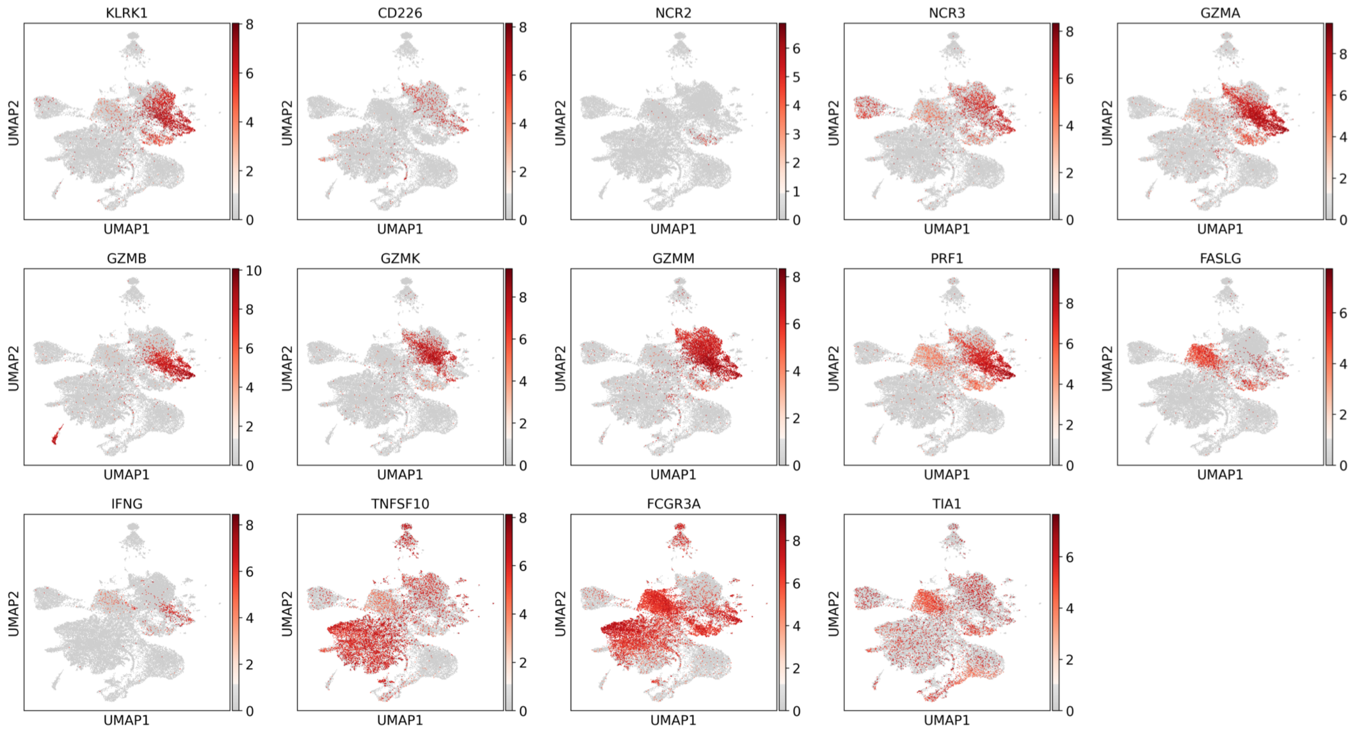

**Figure S6. Cell Cycle phase distribution in Tumor sub-clusters**

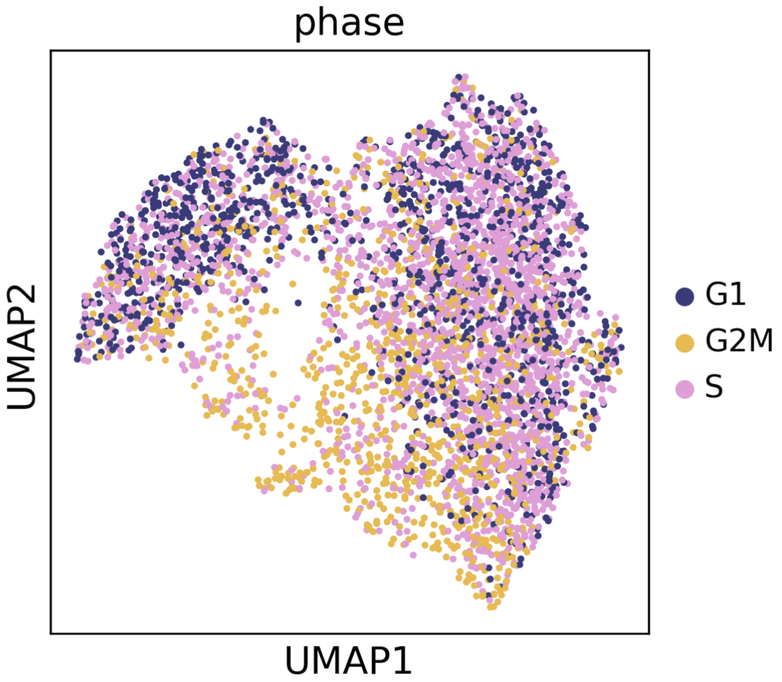

**Figure S7. CD19 was highly expressed in Tumor_1 but not Tumor_2. Other B cell marker were not expressed in both Tumor sub-clusters**

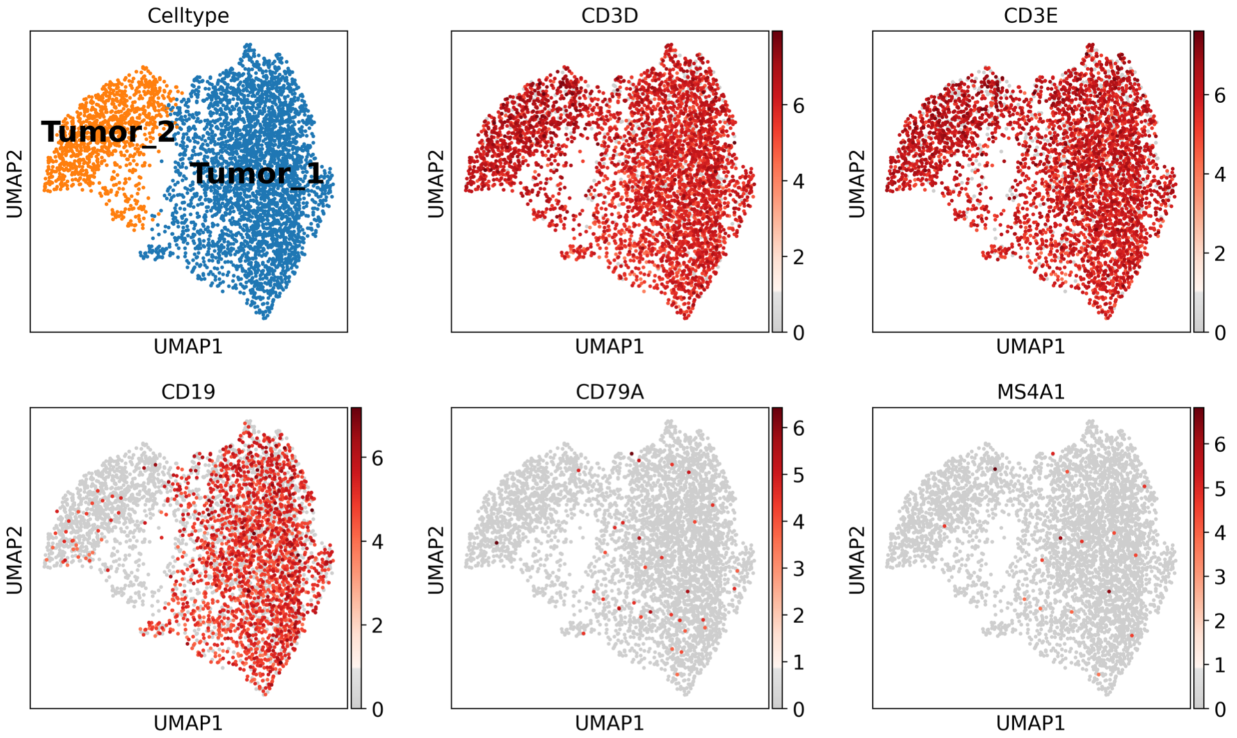

**Figure S8. T cell sub-clusters identified from the tumor microenvironment.** (A and B) The expression of marker genes for the T cell sub-clusters annotation

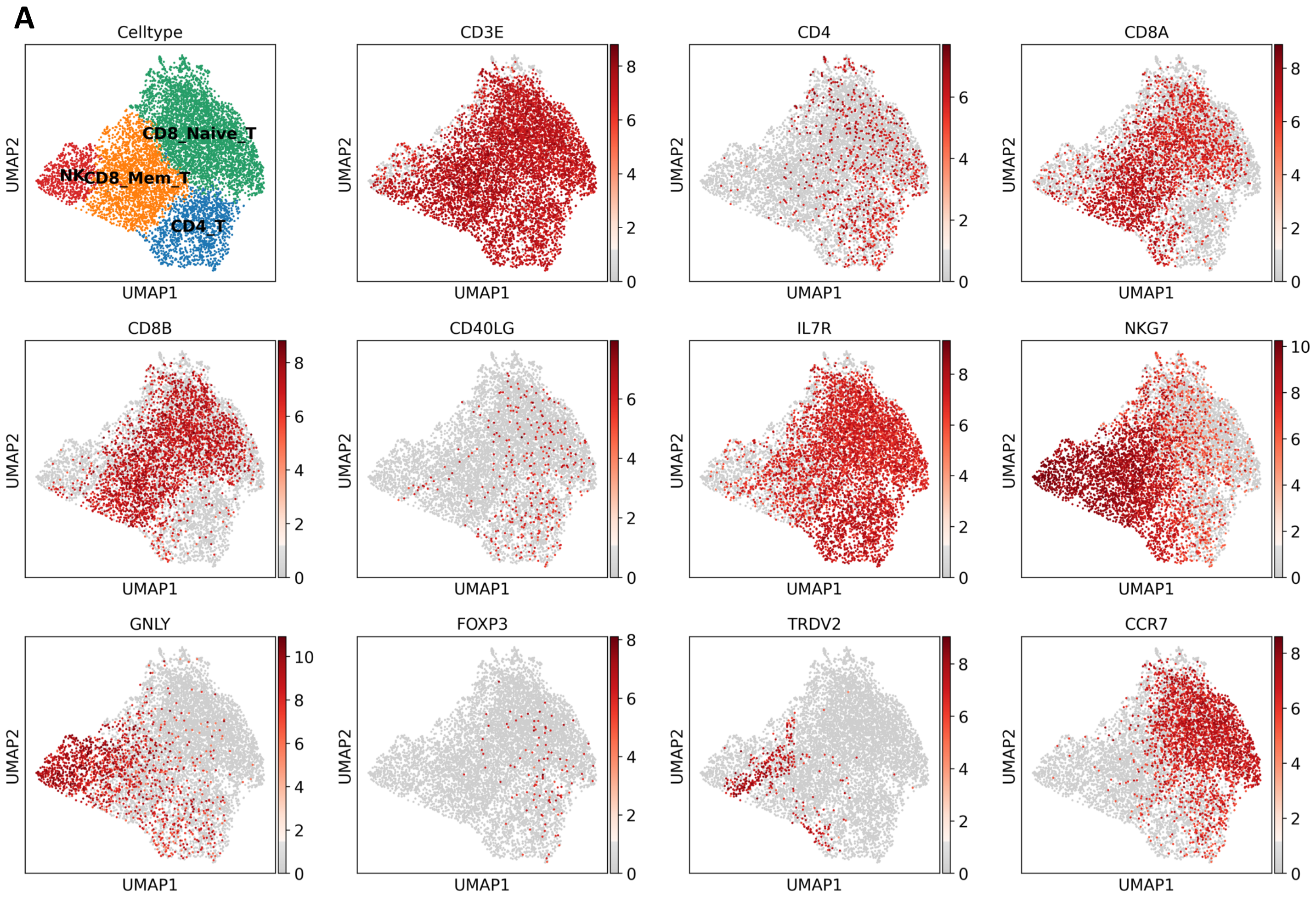


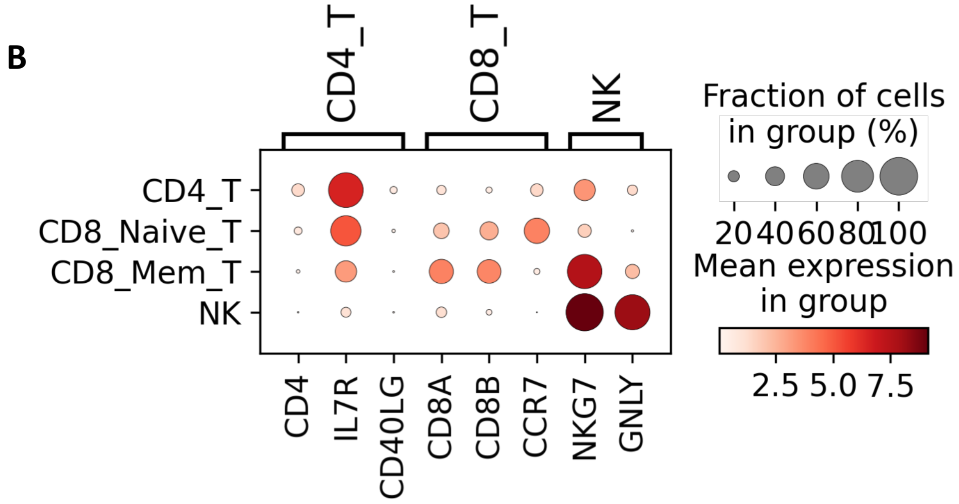

### Figure S9. Total interaction counts and interaction strength inferred from each sample

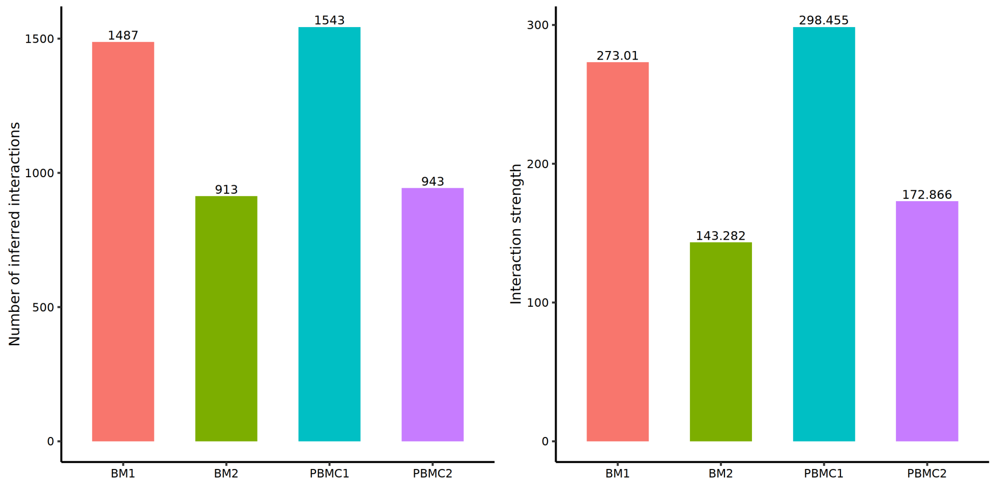
